# Multidrug treatment with nelfinavir and cepharanthine against COVID-19

**DOI:** 10.1101/2020.04.14.039925

**Authors:** Hirofumi Ohashi, Koichi Watashi, Wakana Saso, Kaho Shionoya, Shoya Iwanami, Takatsugu Hirokawa, Tsuyoshi Shirai, Shigehiko Kanaya, Yusuke Ito, Kwang Su Kim, Kazane Nishioka, Shuji Ando, Keisuke Ejima, Yoshiki Koizumi, Tomohiro Tanaka, Shin Aoki, Kouji Kuramochi, Tadaki Suzuki, Katsumi Maenaka, Tetsuro Matano, Masamichi Muramatsu, Masayuki Saijo, Kazuyuki Aihara, Shingo Iwami, Makoto Takeda, Jane A. McKeating, Takaji Wakita

## Abstract

Antiviral treatments targeting the emerging coronavirus disease 2019 (COVID-19) are urgently required. We screened a panel of already-approved drugs in a cell culture model of severe acute respiratory syndrome coronavirus 2 (SARS-CoV-2) and identified two new antiviral agents: the HIV protease inhibitor Nelfinavir and the anti-inflammatory drug Cepharanthine. *In silico* modeling shows Nelfinavir binds the SARS-CoV-2 main protease consistent with its inhibition of viral replication, whilst Cepharanthine inhibits viral attachment and entry into cells. Consistent with their different modes of action, *in vitro* assays highlight a synergistic effect of this combined treatment to limit SARS-CoV-2 proliferation. Mathematical modeling in vitro antiviral activity coupled with the known pharmacokinetics for these drugs predicts that Nelfinavir will facilitate viral clearance. Combining Nelfinavir/Cepharanthine enhanced their predicted efficacy to control viral proliferation, to ameliorate both the progression of disease and risk of transmission. In summary, this study identifies a new multidrug combination treatment for COVID-19.

## Introduction

The novel coronavirus infectious disease 2019 (COVID-19), caused by the severe acute respiratory syndrome coronavirus 2 (SARS-CoV-2), is a global public health problem that is impacting social and economic damage worldwide (Huang et al., 2020; Zhou et al., 2020; Zhu et al., 2020). As of April 12, 2020, 1,696,588 confirmed cases with 105,952 deaths were reported across 213 countries/areas/territories (WHO). COVID-19 was characterized as a pandemic by the World Health Organization (WHO), however, there is no approved treatment. Several drugs have been evaluated in COVID-19 patients in clinical trials: including Lopinavir (LPV) and Ritonavir, Chloroquine (CLQ), Favipiravir (FPV), and Interferon, all repurposed FDA-approved drugs, together with Remdesivir (RDV), an antiviral agent that awaits clinical approval (Cao et al., 2020; Dong et al., 2020; Touret and de Lamballerie, 2020). The clinical efficacies of these drugs are expected shortly, however, additional treatment options are urgently needed.

In this study, we screened a panel of FDA/EMA/PMDA-approved drugs in a SARS-CoV-2 infection cell culture assay and identified two, Nelfinavir (NFV) and Cepharanthine (CEP), that show more potent antiviral activity in this *in vitro* screen compared to drugs currently being trialed. Our screen shows that both NFV and CEP inhibit SARS-CoV-2 at concentrations that can be achieved in the clinic and their different modes of action provide an exciting opportunity for combined multidrug treatment against COVID-19.

## Results

### Anti-SARS-CoV-2 activity of Nelfinavir and Cepharanthine

We established a cell-based drug screening system to identify compounds that protect cells from SARS-CoV-2-induced cytopathology (Fig. 1A): VeroE6/TMPRSS2 cells were treated with compounds for 1 h during inoculation with a clinical isolate of SARS-CoV-2 (Matsuyama et al., 2020) at a multiplicity of infection (MOI) of 0.01. Unbound virus was removed by washing and the cells treated with compounds for 48 h to assess cell viability (Fig. 1A) (Methods). SARS-CoV-2 replication in VeroE6/TMPRSS2 induced a cytopathic effect and to validate our assay we show that two compounds, LPV and CLQ, that were reported to inhibit SARS-CoV-2 infection (Wang, M. et al., 2020), reduced virus-induced cytopathicity (Fig. 1B, compare b and c, d).

**Fig. 1.**
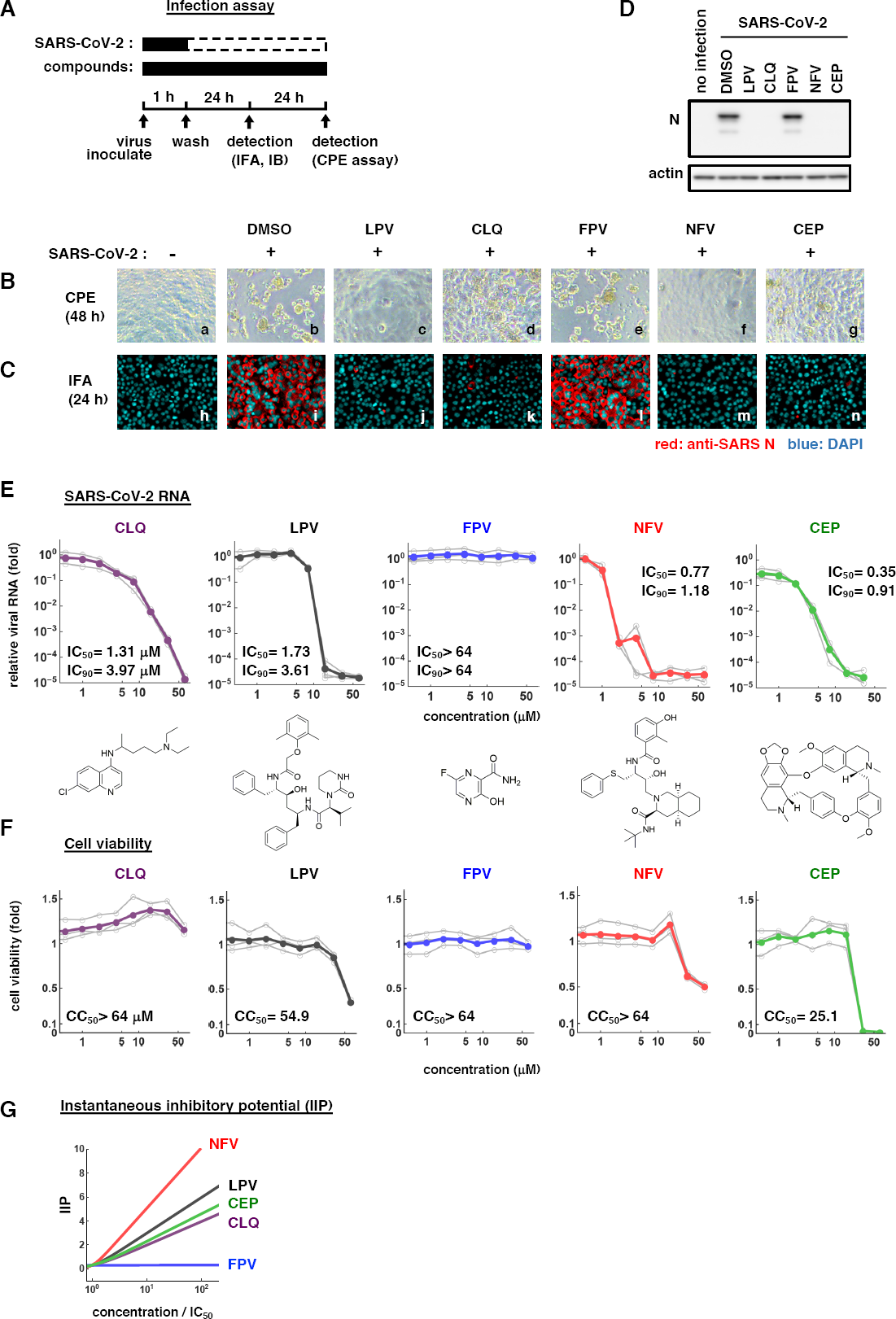
Nelfinavir (NFV) and Cepharanthine (CEP) inhibit SARS-CoV-2 infection. **(A)** Schematic of the SARS-CoV-2 infection assay. VeroE6/TMPRSS2 cells were inoculated with SARS-CoV-2 at an MOI=0.01 in the presence of compounds. After washing out unbound virus, the cells were incubated with compounds for 24-48 h. Cells were harvested for immunofluorescence (IFA) or immunoblot analyses of viral N protein at 24 h and cytopathic effects (CPE) at 48 h post-infection. Solid and dashed boxes indicate the periods with and without treatment, respectively. **(B)** Virus-induced CPE following drug treatment was recorded at 48 h post-infection. Immunofluorescence **(C)** and immunoblot **(D)** detection of viral N protein expression in the infected cells at 24 h post-infection, where the red and blue signals show N and DAPI, respectively. Dimethyl sulfoxide (DMSO), 0.4%; Lopinavir (LPV), 16 μM; Chloroquine (CLQ), 16 μM; Favipiravir (FPV), 32 μM; NFV, 4 μM; CEP, 8 μM. **(E, F)** Dose-response curves for compounds. In (E), secreted viral RNA at 24 h post-inoculation was quantified and plotted against drug concentration and chemical structures shown below each graph (for CEP, the structure of a major component is shown). In (F), viability of cells treated with the compounds was quantified by MTT assay. Inferred IC_50_, IC_90_, and CC_50_ values are shown. **(G)** The antiviral activity for each drug is determined and Instantaneous inhibitory potential (IIP) shown.

After screening 306 FDA/EMA/PMDA-approved drugs, we identified compounds that protected cell viability by 20-fold compared with a DMSO solvent control (Methods) (Supplementary Table S1). Among these, we selected to study NFV and CEP as candidates showing the greatest anti-cytopathic activity (Fig. 1B, f and g). NFV targets human immunodeficiency virus (HIV) protease and CEP is a Stephania-derived alkaloid extract having anti-inflammatory and anti-oxidative activities (Bailly, 2019; Kao et al., 2015; Markowitz et al., 1998). To confirm and extend these observations we assessed SARS-CoV-2 encoded N protein expression 24 h post-inoculation by immunofluorescence (Fig. 1C, red) and immunoblotting (Fig. 1D). Both NFV and CEP significantly reduced N protein expression, confirming these compounds inhibit SARS-CoV-2 proliferation. To quantify their anti-SARS-CoV-2 activity, we treated cells with a range of drug concentrations and measured secreted viral RNA 24 h post-infection. NFV and CEP, together with CLQ and LPV, significantly reduced viral RNA levels in a dose-dependent manner to 0.001 ∼ 0.01% of the untreated control infections (Fig. 1E). FPV showed negligible antiviral activity against SARS-CoV-2, consistent with previous reports (Choy et al., 2020; Wang, M. et al., 2020). In parallel we assessed cell viability and noted cell death at high drug concentrations up to 64 μM (Fig. 1F). The concentration of drugs required to inhibit 50% (IC_50_) or 90% (IC_90_) of virus replication along with their 50% cytotoxicity (CC_50_) are listed in Fig. 1E and F. These experiments highlight a > 70-fold window (CC_50_/IC_50_) where NFV and CEP can inhibit SARS-CoV-2 proliferation with minimal toxicity.

We previously reported a method to quantify instantaneous inhibitory potential (IIP) (Koizumi et al., 2017) and imply that NFV and CEP will have higher antiviral potentials than LPV and CLQ, respectively (Supplementary Note, Supplementary Table S2).

### Modes of action of Nelfinavir and Cepharanthine

To define how these compounds impact on the viral replicative life cycle, we performed a time of addition assay (Fig. 2A). We measured the antiviral activity of drugs added at different times: (a) present during the 1 h virus inoculation step and maintained throughout the 24 h infection period (**whole life cycle**); (b) present during the 1 h virus inoculation step and for an additional 2 h and then removed (**entry**); or (c) added after the inoculation step and present for the remaining 22 h of infection (**post-entry**). CLQ, a known modulator of intracellular pH that non-specifically inhibits virus entry (Akpovwa, 2016), was recently reported to inhibit SARS-CoV-2 (Liu, J. et al., 2020; Wang, M. et al., 2020) and we confirmed its activity in the early stages of infection (Fig. 2B, lane 8). RDV was previously reported to inhibit the process for intracellular viral replication (Wang, M. et al., 2020) and we confirmed this mode of action showing a reduction in viral RNA levels with a negligible effect on virus entry (Fig. 2B, lane 5).

**Fig. 2.**
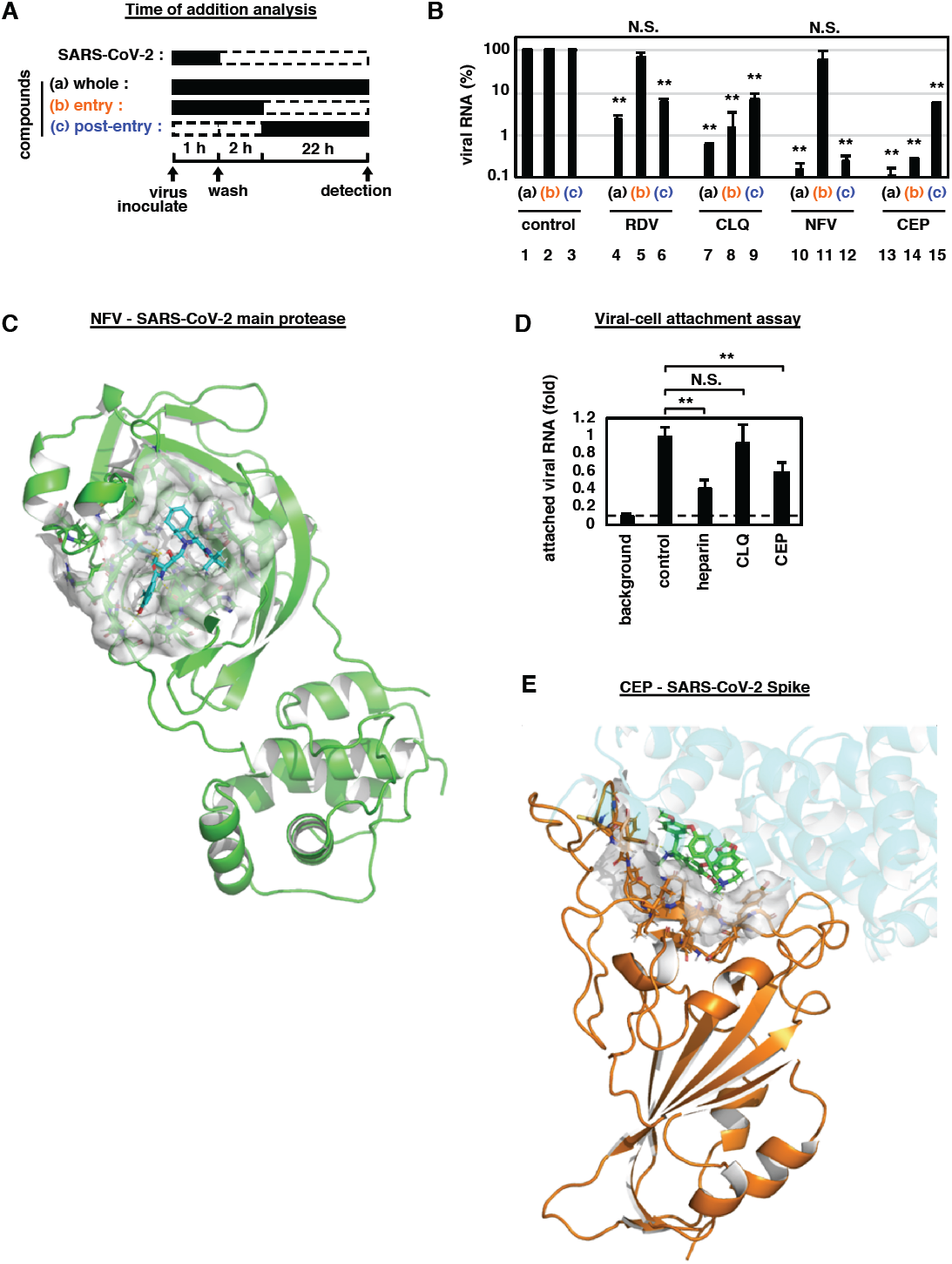
Antiviral modes of action for NFV and CEP. **(A, B)** Time of addition analysis to examine steps in SARS-CoV-2 life cycle. (A) shows the schematic of the time of addition analysis. Compounds were added at different times (a, whole; b, entry; or c, post-entry): (a): presentation during the 1h virus inoculation step and maintained throughout the 24 h infection period (**whole life cycle**); (b) present during the 1 h virus inoculation step and for an additional 2 h and then removed (**entry**); or (c): added after the inoculation step and present for the remaining 22 h of infection (**post-entry**). Solid and dashed boxes indicate the periods with and without treatment, respectively. In (B), the antiviral activities of each compound under the various protocols are estimated by quantifying the levels of secreted viral RNA at 24 h post-inoculation. **(C)** Predicted binding of NFV to SARS-CoV-2 main protease. Representation of SARS-CoV-2 main protease (green), NFV molecule (cyan stick) and protease binding site residues around NFV within 4 Å (surface representation) are shown. **(D**) Virus-cell attachment assay. VeroE6/TMPRSS2 cells were incubated with virus (MOI=0.001) in the presence of the indicated compounds for 5 min at 4°C to allow virus-cell attachment with no internalization. After extensive washing, viral RNA on the cell surface was quantified, where the background depicts residual viral inocula in the absence of cells. **(E)** Predicted binding of CEP molecule to SARS-CoV-2 Spike protein. Spike protein, CEP molecule and protein binding site residues around CEP within 4 Å are shown in cartoon representation colored in orange, green stick and surface representation, respectively. An Overlapping view of the ACE2 with CEP is shown in semi-transparent cartoon representation colored in cyan.

This assay identified NFV as a replication inhibitor whilst CEP targeted the virus entry phase (Fig. 2B, lanes 10-15). These data are consistent with reports that LPV and NFV inhibited the replication of other coronaviruses, SARS-CoV (Liu et al., 2005; Wu et al., 2004; Yamamoto et al., 2004), and that CEP reduced the entry of human coronavirus OC43 (Kim et al., 2019).

Since NFV binds the HIV-1 protease we used an *in silico* docking simulation to assess its potential interaction with the SARS-CoV-2 encoded main protease (Fig. 2C). NFV was ranked in the top 1.5% of compounds following an *in silico* screen of the SARS-CoV-2 encoded main protease (see Methods) (Fig. 2C, cyan stick: NFV, green: main protease). Our docking model predicts that NFV interacts with the SARS-CoV-2 protease active site pocket and would block the recruitment of substrates.

To investigate whether CEP inhibits SARS-CoV-2 particle attachment or internalization into cells, we established an assay to measure viral attachment to cells by pre-chilling to inhibit particle endocytosis. Cell-bound virus particles are measured by qPCR of viral RNA. Viruses frequently exploit cellular heparan sulfate proteoglycans to initiate low affinity attachment and heparin shows broad-spectrum inhibition of virus-cell attachment (De Clercq, 1998; Lang et al., 2011). Unsurprisingly, heparin blocked SARS-CoV-2 particle attachment to VeroE6/TMPRSS2 cells (Fig. 2D). We demonstrate that CEP significantly inhibited SARS-CoV-2 attachment to cells, whereas CLQ that targets intracellular trafficking pathways (Liu, J. et al., 2020) had no effect (Fig. 2D). *In silico* docking simulation confirms that CEP molecule (a major component of the pharmaceutical preparation of CEP) can bind to SARS-CoV-2 Spike protein and interfere with the Spike engagement to its receptor, angiotensin-converting enzyme 2 (ACE2) (Lan et al., 2020; Walls et al., 2020; Wang, Q. et al., 2020) (Fig. 2E, green stick: CEP molecule, orange: Spike, semi-transparent cyan: ACE2). These data highlight a new role for CEP to inhibit SARS-CoV-2 particle attachment to cells.

### Nelfinavir and Cepharanthine show synergistic antiviral activity

NFV showed a modest increase in antiviral activity (IC_90_ of 1.18 μM) compared to LPV (IC_90_ of 3.61 μM), similarly CEP (IC_90_ of 0.91 μM) showed greater antiviral activity than CLQ (IC_90_ of 3.97 μM). Importantly, both NFV and CEP show anti-SARS-CoV-2 activity within the concentration ranges achieved in patients, where the C_max_ of both drugs are 6.9 and 2.3 μM (by administration of 500 mg NFV orally and of 100 mg CEP by intravenous injection) respectively (Markowitz et al., 1998; Yasuda et al., 1989). Since NFV and CEP have different mode of actions, we examined their potential for synergistic effects. Antiviral activity and cell viability were determined by qPCR enumeration of viral RNA and MTT activity, respectively, following treatment with each compound alone or in combination (Fig. 3). For these experiments, we infected cells with lower amounts of SARS-CoV-2 (MOI=0.001) than used in our earlier drug screen. Single treatment with NFV or CEP reduced viral RNA in a dose-dependent manner and co-treatment further reduced viral RNA levels (Fig. 3A): e.g. NFV (2.24 μM) or CEP (3.20 μM) alone reduced viral RNA to 5.8% and 6.3% of untreated control, respectively, however, their combination reduced viral RNA level to 0.068%. Higher doses of the NFV/CEP combined treatment (4 μM each) reduced the viral RNA to undetectable levels. We compared the observed experimental antiviral activity (Fig. 3A, Supplementary Fig. S1A) with theoretical predictions calculated using a classical Bliss independence method that assumes the drugs act independently (Supplementary Note, Supplementary Fig. S1B) (Greco et al., 1995; Koizumi and Iwami, 2014). The difference between the observed values and theoretical predictions suggest that NFV and CEP exhibit a synergistic activity over a broad range of concentrations (Fig. 3C red: synergistic effect).

**Fig. 3.**
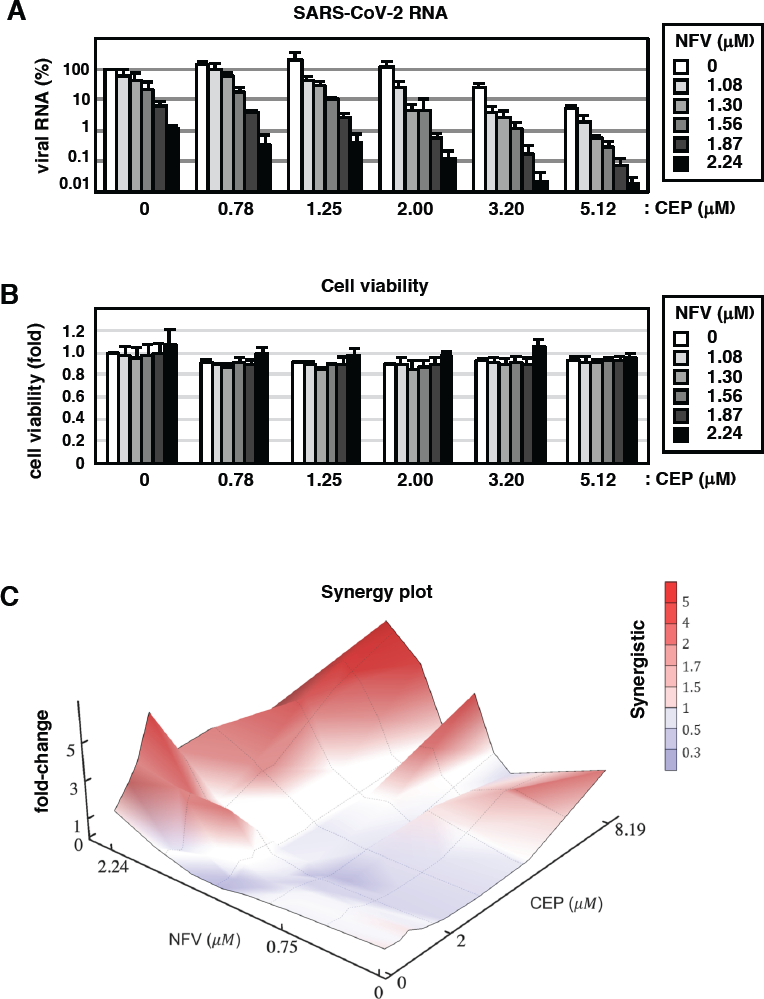
Combination treatment with NFV and CEP. **(A)** Dose-response curve of NFV/CEP co-treatment in the infection experiment (MOI=0.001). Extracellular viral RNA levels at 24 h post-infection were quantified and plotted against concentrations of NFV (1.08, 1.30, 1.56, 1.87, and 2.24 μM) and CEP (0.78, 1.25, 2.00, 3.20, and 5.12 μM). **(B)** Cell viability upon co-treatment with compounds. **(C)** The three-dimensional interaction landscapes of NFV and CEP were evaluated based on the Bliss independence. Red and blue colors on the contour plot indicate synergy and antagonism, respectively.

### Modeling the impact of Nelfinavir and Cepharanthine on SARS-CoV-2 dynamics

Combining the published clinical pharmacokinetics information for these drugs with our observed dose-dependent antiviral activities, we can predict the time-dependent antiviral activity (Fig. 4A: left, NFV oral; center, CEP intravenous drip; right: CEP oral) and the resultant viral load dynamics after drug administration (Fig. 4B, Supplementary Note, Supplementary Fig. S2). From such viral dynamics shown in Fig. 4B, we calculated the cumulative viral RNA burden (i.e., area under the curve of viral load) (Fig. 4C, upper) and the time period to reduce the viral load to undetectable levels (Fig. 4C, lower). Our modeling analysis predict that NFV would reduce the cumulative viral load by 91.4% (Fig. 4C, upper, red) and would require 11.9 days to eliminate virus (Fig. 4B, upper left, red), 3.98 days shorter than non-treatment condition (Fig. 4C, lower, red). In contrast, treatment with CEP alone showed a limited effect on the viral load [Fig. 4B, upper right or lower left, green], most likely reflecting the low concentration of the drug when administered orally or by intravenous drip (see Discussion). However, co-administering NFV (oral) and CEP (intravenous drip) resulted in a more rapid decline in viral RNA, with undetectable levels 5.5 days earlier than non-treatment and 1.5 days earlier than NFV alone (Fig. 4C). Another advantage of combination treatment is discussed in Discussion. In summary, NFV is likely to show antiviral activity at clinically achievable drug concentration and combination treatment with CEP will facilitate virus elimination.

**Fig. 4.**
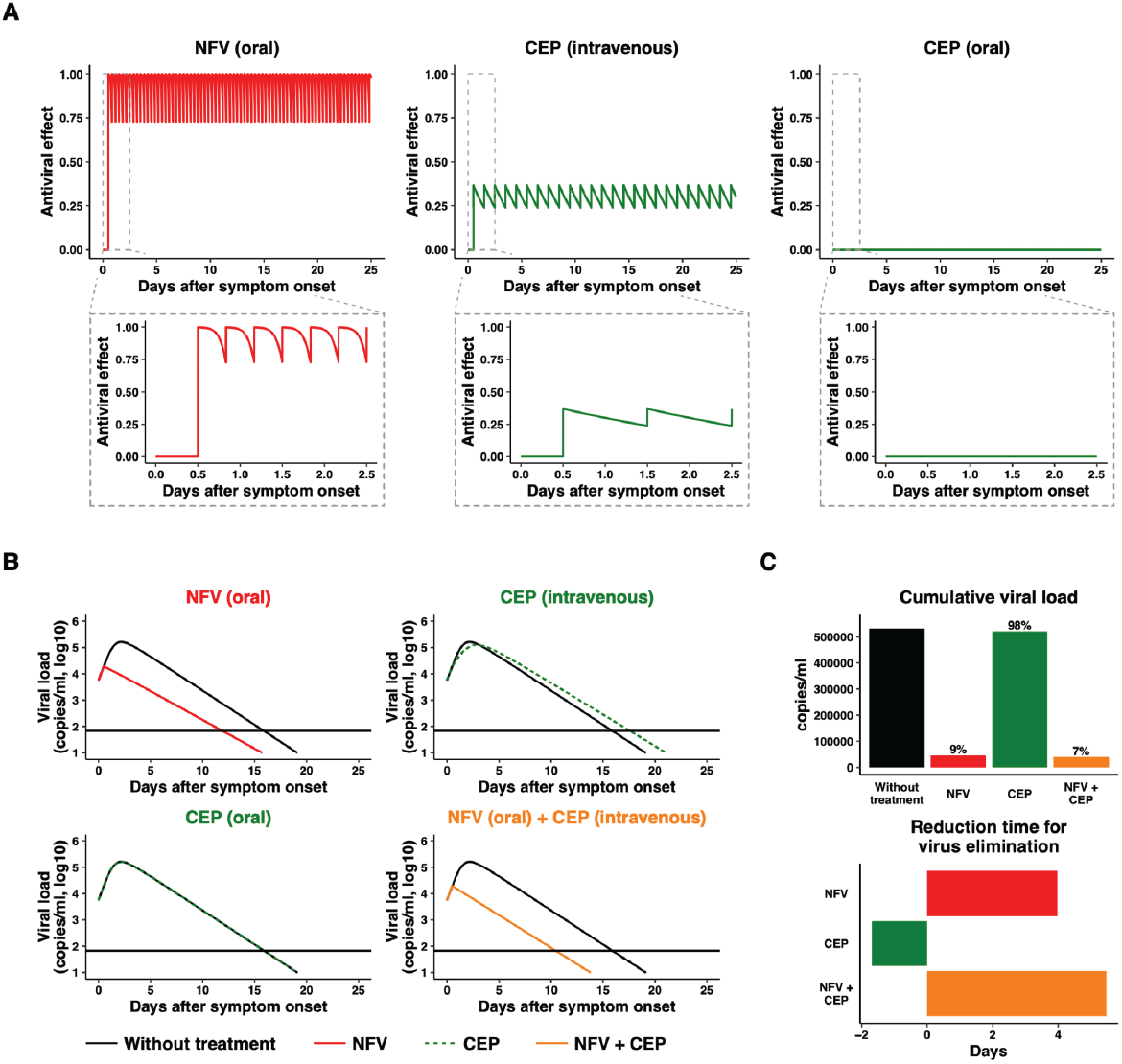
Mathematical prediction of the impact of NFV and CEP therapy on viral dynamics. **(A)** The time-dependent antiviral effects of NFV (500 mg, TID, oral) and CEP [100 mg, intravenous drip or 120 mg, oral] predicted by pharmacokinetics/pharmacodynamics (PK/PD) model are shown with enlarged views of the gray zones in upper panels. **(B)** Viral load dynamics in the presence or absence of NFV (oral), CEP (intravenous), CEP (oral), and NFV (oral)/CEP (intravenous) combined therapies predicted by pharmacokinetics/pharmacodynamics/viral-dynamics (PK/PD/VD) models are shown. **(C)** The cumulative antiviral load [area under the curve in (B)] (upper) and the reduction time (days) for virus elimination (lower) with drug single or combined treatments are shown.

## Discussion

Screening a panel of approved drugs identified two agents, NFV and CEP, with potent antiviral activity against SARS-CoV-2. NFV inhibits SARS-CoV-2 replication and our modeling data suggests this is mediated via a direct interaction with the viral encoded main protease (Fig. 2B and C). A recent study reported that CEP exhibited anti-SARS-CoV-2 activity (Fan et al., 2020), these authors speculated that CEP targeted both the entry and viral replication phase. However, our time of addition experiments suggest that CEP predominantly inhibits viral entry (Fig. 2B, lane 14). Furthermore, virus-cell attachment assays and docking simulations confirm that CEP inhibits virus attachment to target cells (Fig. 2D and E). There is a significant global effort to generate a COVID-19 vaccine that will target the SARS-CoV-2 encoded Spike glycoprotein (Thanh Le et al., 2020) that is required for particle engagement of the receptor ACE2 for infecting cells. We predict that CEP may work synergistically with vaccine induced anti-S antibody responses and such experiments are worthy of future investigation. Further mechanistic studies will be required to confirm the proposed mechanisms of action of these compounds. However, our observation that NFV and CEP target different steps in the viral life cycle support the development of multidrug combination therapies for treating COVID-19.

Our mathematical modeling studies assess how anti-SARS-CoV-2 drug candidates can suppress virus proliferation and facilitate virus elimination (Fig. 4). At clinical doses NFV can maintain strong antiviral effect over time and thus can reduce SARS-CoV-2 RNA burden that results in shortening the time required to eliminate infection. In contrast, CEP monotherapy is predicted to have a modest antiviral effect because of a low concentration *in vivo* when administered by oral or intravenous drip. However, higher doses of CEP, based on its relatively safe toxicity profile (Rogosnitzky and Danks, 2011), may increase drug efficacy in a clinical setting. It is noteworthy that combining CEP with NFV further reduced the cumulative viral load and facilitated virus elimination. As the cumulative viral load in patients is likely to be closely related with the progression of disease and the risk for new transmission (Liu, Y. et al., 2020), such multidrug treatment will be of benefit to improve clinical outcome and to control epidemic. In addition to potentiating antiviral effects, multidrug treatment can limit the emergence of viral drug-resistance which is frequently reported for RNA viruses such as coronavirus.

One limitation of our modeling of drug efficacy is the use of *in vitro* data derived cell culture infection systems without confirmation using *in vivo* infection models. Recently, a SARS-CoV-2 infection system was reported using ferrets, but as yet there is no evidence on the usefulness of this model for evaluating anti-SARS-CoV-2 drugs (Kim et al., 2020). Given the urgency of the problem, this lack of in vivo testing should not prevent the assessment of new antiviral agents. Our screening of approved drugs has identified NFV and CEP as potential anti-SARS-CoV-2 agents. As both NFV and CEP show superior antiviral activities compared to many current drug candidates, these agents offer a promising new multidrug treatment to combat COVID-19.

## Acknowledgments

We thank Drs. Shuetsu Fukushi and Souichi Yamada at Department of Virology I, National Institute of Infectious Diseases for technical assistance. NFV, LPV, and FPV were kindly provided by Japan Tobacco, Abbvie, and Fujifilm Toyama Chemical. Pharmaceutical preparation of CEP was kindly provided by Medisa Shinyaku Inc, a subsidiary of Sawai Pharmaceutical. This work was supported by The Agency for Medical Research and Development (AMED) emerging/re-emerging infectious diseases project (JP19fk0108111, JP19fk0108110, JP20fk0108104); the AMED Basis for Supporting Innovative Drug Discovery and Life Science Research (BINDS, JP19am0101114, JP19am0101069, JP19am0101111) program; The Japan Society for the Promotion of Science KAKENHI (JP17H04085, JP20H03499, JP15H05707, 19H04839); The JST MIRAI program; and Wellcome Trust funded Investigator award (200838/Z/16/Z).

## Author Contributions

Conceptualization, K.W.; Investigation, H.O., K.W., W.S., K.S., S.Iwanami, T.H., T.S., S.K., Y.I., K.S.K., K.N., S.Iwami; Methodology and Resources, S.Ando., T.S., K.M., M.S., M.T., T.W.; Analysis, All the authors; Writing and editing, K.W., T.H., K.A., S.Iwami, J.A.M; Funding Acquisition, K.W., M.T.; Supervision, K.W.

## Competing Interests

No interests

## Notes

### Competing Interest Statement

The authors have declared no competing interest.

