## Supplementary material for "Multidrug treatment with nelfinavir and cepharanthine against COVID-19"

**Supplementary Methods**
**Supplementary Note**
**Supplementary Figure S1-S2**
**Supplementary Table S1-S4**
**Supplementary References**

#### Supplementary Methods

**Cell culture.** VeroE6/TMPRSS2 cells [VeroE6 cells overexpressing transmembrane protease, serine 2 (TMPRSS2) (Matsuyama et al., 2020) were cultured in Dulbecco's modified Eagle's medium (DMEM; Life Technologies) supplemented with 10% fetal bovine serum (FBS; Cell Culture Bioscience), 10 units/mL penicillin, 10 mg/mL streptomycin, 10 mM HEPES (pH 7.4), and 1 mg/mL G418 (Nacalai) at 37˚C in 5% $CO_2$. During the infection assay, 10% FBS was replaced by 2% FBS and G418 removed.

**Reagents.** All the reagents were purchased from Selleck, Enzo Life Sciences, Cayman Chemical, Sigma, MedChemExpress, TCI or kindly donated by pharmaceutical companies (Abbvie, Alps Pharmaceutical, Asahi Kasei Pharma, Astellas Pharma, Bayer, Boehringer Ingelheim, Bristol-Myers Squibb, Chugai Pharmaceutical, Daiichi Sankyo, EA Pharma, Fujifilm Toyama Chemical, Japan Tobacco, Kakenshoyaku, Kissei Pharmaceutical, Kowa, Kyorin Pharmaceutical, Kyowa Pharmaceutical Industry, Maruho, Mitsubishi Tanabe Pharma, Mochida Pharmaceutical, Novartis, Sanofi, SBI Pharmaceuticals, Shionogi, Sumitomo Dainippon Pharma, Sun Pharma, Takeda Pharmaceutical, Teva Takeda Pharma). Note that throughout in this study we used the pharmaceutical preparation of Cepharanthine (kindly provided by Medisa Shinyaku Inc, a subsidiary of Sawai Pharmaceutical), which is a Stephania-derived alkaloid extract containing 19.5-33.5% Cepharanthine molecule as the major component.

**Infection assay.** SARS-CoV-2 was handled in a biosafety level 3 (BSL3). We used the SARS-CoV-2 Wk-521 strain, a clinical isolate from a COVID-19 patient, and obtained viral stocks by infecting VeroE6/TMPRSS2 cells (Matsuyama et al., 2020). Virus infectious titers were measured by inoculating cells with a 10-fold serial dilution of virus and cytopathology measured to calculate $TCID_{50}$/ml (Matsuyama et al., 2020). For the infection assay, VeroE6/TMPRSS2 cells were inoculated with virus at an MOI of 0.01 (Fig. 1 and 2B) or 0.001 (Fig. 2D and 3) for 1 h and unbound virus removed by washing. Cells were cultured for 24 h prior to measuring extracellular viral RNA or detecting viral encoded N protein, and cytopathic effects (CPE) after 48 h. Compounds were added during virus inoculation (1 h) and replenished after washing (24 or 48 h) except for time of addition assay.

For the time of addition assay, we added compounds with three different timings (Fig. 2A): (a) present during the 1 h virus inoculation step and maintained throughout the 24 h infection period (**whole life cycle**); (b) present during the 1 h virus inoculation step and for an additional 2 h and then removed (**entry**); or (c) added after the inoculation step and present for the remaining 22 h of infection (**post-entry**). Inhibitors of viral replication are expected to show antiviral activity in (a) and (c), but not (b), while entry inhibitors (e.g. chloroquine) reduce viral RNA in all three conditions (In c, addition of entry inhibitors after inoculation inhibits re-infection and thus decreases viral RNA) (Wang et al., 2020).

**Quantification of viral RNA.** Viral RNA was extracted with a QIAamp Viral RNA mini kit (QIAGEN) and quantified by real time RT-PCR analysis with a one-step qRT-PCR kit (THUNDERBIRD Probe One-step qRT-PCR kit, TOYOBO) using 5'- ACAGGTACGTTAATAGTTAATAGCGT-3' and 5'-

ATATTGCAGCAGTACGCACACA-3’ as the primer set and a 5’-FAM-ACACTAGCCATCCTTACTGCGCTTCG-TAMRA-3’ probe, as described (Corman et al., 2020). Detection limit of SARS-CoV-2 RNA in this study was 38 cycle as $C_t$ cycle.

**Detection of viral N protein.** Viral protein expression was detected using a rabbit anti-SARS-CoV N antibody (Mizutani et al., 2004) with AlexaFluor 568 anti-rabbit IgG or anti-rabbit IgG-HRP (Thermo Fisher) by indirect immunofluorescence or immunoblot analyses as previously reported (Ohashi et al., 2018).

**Cell viability and virus induced cytopathology.** Cell viability was determined by MTT assay as previously reported (Ohashi et al., 2018). Virus-induced cytopathology was observed by microscopy at 48 h post-infection as previously reported (Matsuyama et al., 2020).

**Chemical screening.** We screened an FDA/EMA/PMDA-approved chemical library composed of 306 compounds. Cells were treated with compounds at 8, 16, or 30 μM for 1 h during virus inoculation and for up to 72 h post-inoculation. The cells were then fixed with 4% paraformaldehyde and stained with DAPI to count viable cells using a high-content imaging system (ImageXpress Micro Confocal, Molecular Devices). Compounds that protected cells from virus-induced cytopathology and showed survival cell number more than 20-fold of the control were selected as hits. The list of compounds used in this study is shown in Table S1. Among 306 tested compounds, Cepharanthine, Lopinavir, Loteprednol, Nelfinavir, and Rapamycin were selected as hits. Lopinavir is already underway for clinical trial as anti-SARS-CoV-2 agents (Cao et al., 2020). As Loteprednol and Rapamycin are steroid and immunosuppressant, respectively, we focused on Cepharanthine and Nelfinavir in this study.

**Docking simulation of a target protein and a compound.** The crystal structure of the main protease and spike protein were obtained from Protein Data Bank (6LU7 (Jin et al., 2020) and 6M0J (Lan et al., 2020)) and refined for docking simulations using the Protein Preparation Wizard Script within Maestro (Schrödinger, LLC). We carried out *in silico* library screening based on the active site pocket of the main protease using combined molecular docking with a protein-ligand interaction fingerprint scoring method against 8,085 known drugs obtained from the KEGG-Drug database (Kanehisa and Goto, 2000). For all compounds ionization and energy minimization were performed by the OPLS3 force field in the LigPrep Script of Maestro (Schrödinger, LLC). These minimized structures were used as input structures for docking simulations. Docking simulations were performed using the Glide (Friesner et al., 2004; Halgren et al., 2004) SP docking program (Schrödinger, LLC) with a grid box defined by N3 inhibitor molecule for main protease and ACE2 binding interface residues for spike protein using BioLuminate (Schrödinger, LLC)

**Mathematical analysis.** Determination of synergism between NFV and CEP and simulation of virus dynamics as well as the calculation of IIP are shown in Supplementary Note in detail.

**Statistics.** All experiments were repeated three times in each assay. Statistical significance estimated using the two-tailed Student's t test (*$p<0.05$; **$p<0.01$; N.S., not significant).

#### Supplementary Note

##### Quantifying instantaneous inhibitory potential (IIP) from the dose-response curves of the drugs

The typical dose-response curves of a single antiviral drug can be analyzed using the following Hill function (Koizumi et al., 2017) (**Fig. 1E**):

$$f_u = \frac{1}{1 + \left(\frac{D}{IC_{50}}\right)^m}. \qquad (1)$$

Here, $f_u$ represents the fraction of infection events unaffected by the drug (i.e., $1 - f_u$ equals the fraction of drug-affected events). $D$ is the drug concentration, $IC_{50}$ is the drug concentration that achieves 50% inhibition of activity, and $m$ is the slope of the dose-response curve (i.e., Hill coefficient) (Koizumi et al., 2017). Dose-response curves for drugs with higher $m$ values show stronger antiviral activity at the same normalized drug concentration so long as the drug concentration is higher than $IC_{50}$ (**Fig. 1E**). Least-square regression approach was used to fit Eq.(1) to dose-response data and estimate the values of $IC_{50}$ and $m$. Those estimated values for each drug against SARS-CoV-2 are summarized in **Table S2**.

We evaluated the intrinsic antiviral activity of anti-SARS-CoV-2 agents (**Fig. 1G**). The antiviral activity of antiviral drugs can be expressed as the instantaneous inhibitory potential (IIP) (Jilek et al., 2012; Laskey and Siliciano, 2014; Sampah et al., 2011; Shen et al., 2008; Shen et al., 2011; Shen et al., 2009):

$$\mathrm{IIP} = \log\left(\frac{1}{f_u}\right) = \log\left[1 + \left(\frac{D}{IC_{50}}\right)^m\right]. \qquad (2)$$

If a drug reduces SARS-CoV-2 replication by 1 log then $f_u = 0.1$ and its $\mathrm{IIP} = 1$, whereas if it reduces replication by 2 logs, i.e. 100-fold, its $\mathrm{IIP} = 2$. Note that the IIP incorporates all three parameters of the dose-response curve; $D$, $IC_{50}$ and $m$. Eq. (2) indicates that the higher the $m$ of the drug, the higher the IIP at a given $D$ and $IC_{50}$.

##### Expected anti-SARS-CoV-2 effect of double-drug combinations by Bliss independence

We evaluated the effect of double-drug combinations for Bliss independence which is widely used to analyze drug combination data (Bliss, 1939; Kobayashi et al., 2014; Koizumi and Iwami, 2014; Tallarida, 2001). Bliss independence assumes that each drug acts on different targets, and is defined as:

$$f_u^{\mathrm{Bcom}} = f_u^A(D) \times f_u^B(D), \qquad (3)$$

where $f_u^{\mathrm{Bcom}}$, $f_u^A$ and $f_u^B$ are the fractions of infection events unaffected by the combined drugs A (i.e., Nelfinavir: NFV) and B (i.e., Cepharanthine: CEP) expected by the Bliss model, single drug A and single drug B defined by Eq. (1), respectively. Using Eq. (2), we expected the anti-SARS-CoV2 effects of combined drugs A and B, $1 - f_u^{\mathrm{Bcom}}$, from the anti-SARS-CoV-2 effects of the single drugs (**Fig. S1**).

However, the Bliss model ignores interactions in which drugs enhance each other effects. To address this point, we introduced the recent proposed model (Zimmer et al., 2016), called “dose model” considering the drug interactions, and further evaluated the expected antiviral effects (**Fig. S1**). This drug interaction is described by introducing interaction terms between drug pairs, that is, the “effective” concentration of drug A (i.e., NFV) and B (i.e., CEP), $D_A^{\mathrm{com}}$ and $D_B^{\mathrm{com}}$, are defined as follows;

$$D_A^{\mathrm{com}} = D_A\left(1 + a_{AB}\frac{D_B^{\mathrm{com}}}{IC_{50}^B + D_B^{\mathrm{com}}}\right)^{-1}, \qquad D_B^{\mathrm{com}} = D_B\left(1 + a_{BA}\frac{D_A^{\mathrm{com}}}{IC_{50}^A + D_A^{\mathrm{com}}}\right)^{-1},$$

where $D_A$ and $D_B$ are the “true” concentrations, $IC_{50}^A$ and $IC_{50}^B$ are the concentrations that achieve 50% inhibition of activity, $a_{AB}$ and $a_{BA}$ are the interaction parameters for drug A and B, respectively. Note that $IC_{50}^A$ and $IC_{50}^B$ are corresponding to the estimations from the dose-response curves of a single antiviral drug in **Table S2**, and $a_{AB} = -0.462$ and $a_{BA} = 0.307$ are estimated from the dose-response curves of the double-drug combination. The dose model extended the Bliss model, thus, the expected anti-SARS-CoV-2 effect with effective concentration of drugs A and B (rather than the true concentrations) are calculated as $1 - f_u^{\mathrm{Dcom}}(D)$ and

$$f_u^{\mathrm{Dcom}} = f_u^A(D_A^{\mathrm{com}}) \times f_u^B(D_B^{\mathrm{com}}). \tag{4}$$

The dose model assumed that the effects of drugs on each other’s effective doses are multiplicative.

**PK/PD/VD model for single- and double-drug combinations against SARS-CoV-2 infection**

Based on a standard viral dynamics (VD) model (Ikeda et al., 2016), to describe COVID-19 dissemination among susceptible target cells, we used the following simple mathematical model proposed in (Kim et al., 2020):

$$\frac{df(t)}{dt} = -\beta f(t)V(t), \tag{5}$$

$$\frac{dV(t)}{dt} = \gamma f(t)V(t) - \delta V(t), \tag{6}$$

where $f(t)$ and $V(t)$ are the ratio of uninfected target cells and the amount of virus, respectively. The parameters $\beta$, $\gamma$, and $\delta$ represent the rate constant for virus infection, the maximum rate constant for viral replication and the death rate of infected cells, respectively. All viral load data including Singapore and Zhuhai patients (Young et al., 2020; Zou et al., 2020) were simultaneously fitted using a nonlinear mixed-effect modelling approach, which uses the whole samples to estimate population parameters while accounting for inter-individual variation. The estimated parameters and initial values used here are summarized in **Table S4**.

To investigate the expected outcome for anti-SARS-CoV-2 therapies with single-drug, we conducted *in silico* experiments with the following PK/PD/VD model for replication inhibitor such as Nelfinavir (**Fig. 4**);

$$\frac{df(t)}{dt} = -\beta f(t)V(t), \tag{7}$$

$$\frac{dV(t)}{dt} = \big(1 - \varepsilon(t) \times H(t)\big)\gamma f(t)V(t) - \delta V(t), \tag{8}$$

and for entry inhibitor such as Cepharanthine;

$$\frac{df(t)}{dt} = -\big(1-\eta(t)\times H(t)\big)\beta f(t)V(t), \qquad (9)$$

$$\frac{dV(t)}{dt} = \big(1-\eta(t)\times H(t)\big)\gamma f(t)V(t) - \delta V(t). \qquad (10)$$

Here $H(t)$ is a Heaviside step function defined as $H(t)=0$ if $t<T$: otherwise $H(t)=1$, where $T$ is the initiation timing of the treatment, and the anti-SARS-CoV2 effect for $t>T$ are described as

$$\varepsilon(t)\ (\text{or}\ \eta(t)) = 1 - f_u\big(D(t)\big) = 1 - \frac{1}{1+\left(\frac{D(t)}{IC_{50}}\right)^m}, \qquad D(t) = C_{max}e^{-kt}$$

where $C_{max}$ and $k$ are the peak drug concentration and the elimination rate for corresponding drug, respectively. The parameter values for each drug used here are summarized in **Table S2**.

For anti-SARS-CoV-2 therapies with double-drug combinations, we extended as the following PK/PD/VD model assuming the dose model;

$$\frac{df(t)}{dt} = -\big(1-\eta(t)\times H(t)\big)\beta f(t)V(t), \qquad (11)$$

$$\frac{dV(t)}{dt} = \big(1-\varepsilon(t)\times H(t)\big)\big(1-\eta(t)\times H(t)\big)\gamma f(t)V(t) - \delta V(t). \qquad (12)$$

Here $H(t)$ is a Heaviside step function defined as $H(t)=0$ if $t<T$: otherwise $H(t)=1$, and the anti-SARS-CoV2 effect are described as

$$\varepsilon(t) = 1 - f_u^A\big(D_A^{\text{com}}(t)\ \big) = 1 - \frac{1}{1+\left(\frac{D_A^{\text{com}}(t)}{IC_{50}^A}\right)^{m_A}},$$

$$\eta(t) = 1 - f_u^B\big(D_B^{\text{com}}(t)\big) = 1 - \frac{1}{1+\left(\frac{D_B^{\text{com}}(t)}{IC_{50}^B}\right)^{m_B}},$$

$$D_A^{\text{com}}(t) = C_{\text{max}}^A e^{-k_A t}\left(1 + a_{AB}\frac{D_B^{\text{com}}(t)}{IC_{50}^B + D_B^{\text{com}}(t)}\right)^{-1},$$

$$D_B^{\text{com}}(t) = C_{\text{max}}^B e^{-k_B t}\left(1 + a_{BA}\frac{D_A^{\text{com}}(t)}{IC_{50}^A + D_A^{\text{com}}(t)}\right)^{-1}.$$

Note that we here evaluated the double-drug combination of NFV and CEP (**Fig. 4**), and the pharmacokinetics of NFV and CEP, $D_A^{\text{com}}(t)$ and $D_B^{\text{com}}(t)$, under the combination, are different from those, $D_A(t)$ and $D_B(t)$, under the single-drug treatment because of the effective drug concentration.

**Evaluation of outcomes for anti-SARS-CoV-2 therapies**

The antiviral effect of the anti-viral therapy on SARS-CoV-2 dynamics using Eqs. (7-12) and our estimated parameter values was calculated (**Fig. 4**). We evaluated the outcomes for the therapies defined as "period until virus elimination" and "reduction of cumulative virus production" (**Fig. S2**). Note that the cumulative virus production, i.e., the area under the curve of viral load (AUC: $\int_0^{T_D} V(s)ds$), for SARS-CoV-2 was calculated, where $T_D$ is the time for SARS-CoV-2 achieved the detection limit.

#### Supplementary Figure

##### Fig. S1.

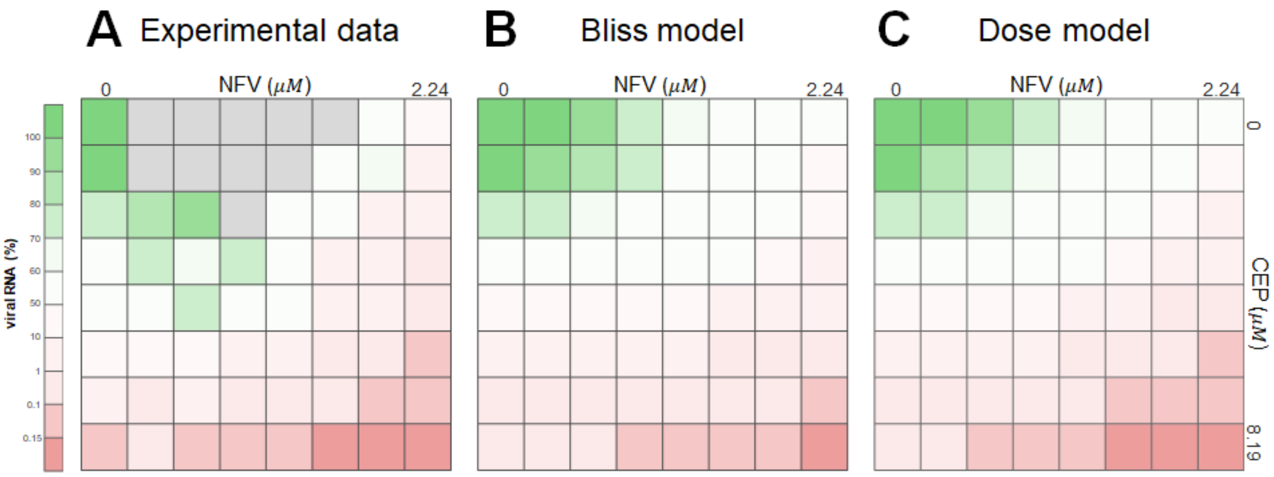


**Fig. S1. Comparison of experimental data, Bliss model and Dose model for the double-drug combinations.** Dose-response matrix of the double-drug combination (corresponding to **Fig. 3A**) are plotted in **(A)**, and the expected anti-SARS-CoV-2 effects of the double-drug combination (NFV and CEP) by the Bliss model and Dose model are plotted in **(B)** and **(C)**, respectively. Note that experimental measurements over 100% of viral RNA (implying large experimental variation because of small dose of antiviral drugs), colored by gray, were excluded in our analysis. The ratios of the values shown in **(A)** over those in **(B)** were calculated and are depicted in **Fig. 3C** in a 3D landscape.

**Fig. S2.**

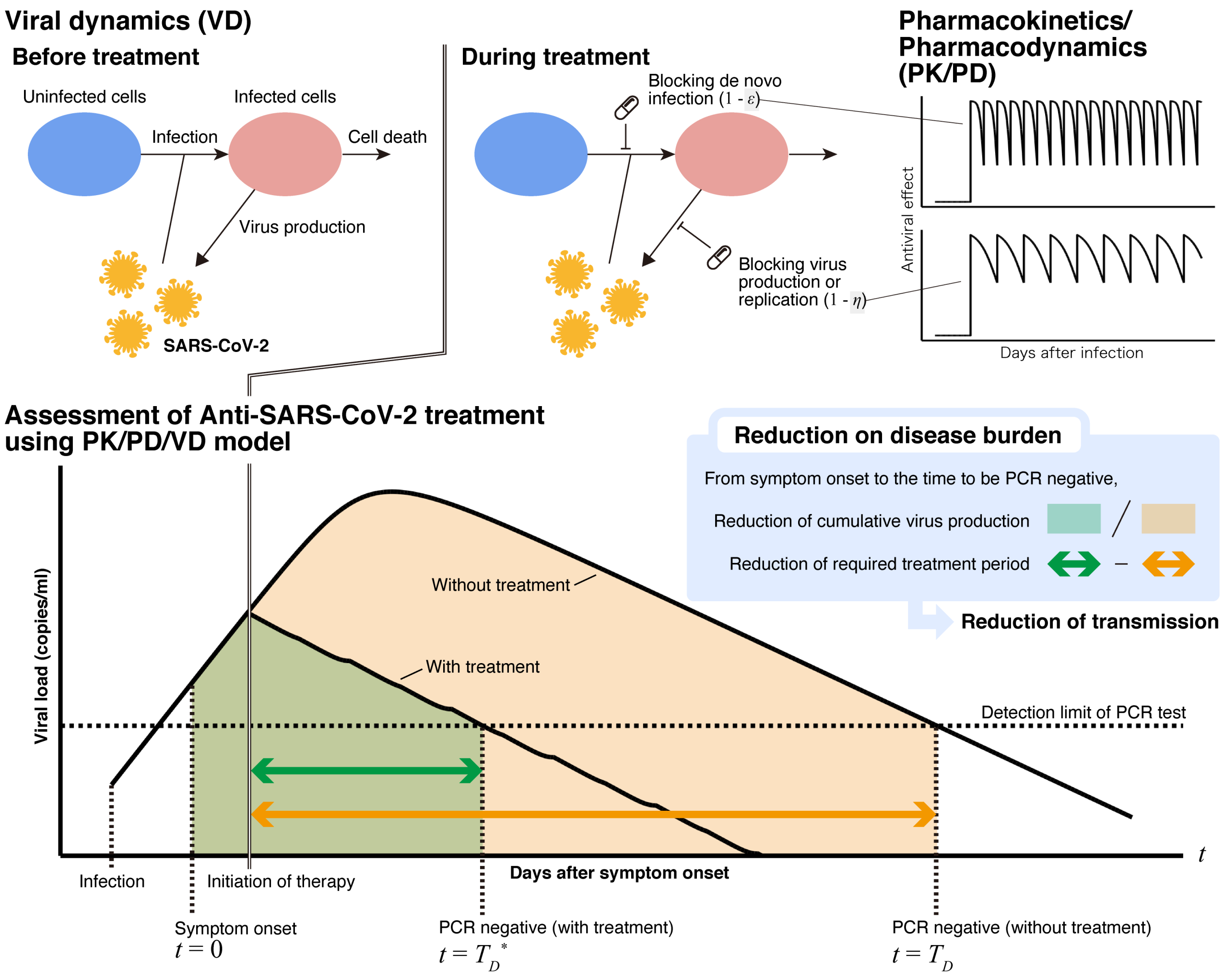


**Fig. S2. Schematic representation of SARS-CoV-2 infection dynamics.** A typical disease progress with viral load on patients undergoing therapy are shown. The outcomes for the therapies, that is, reduction in "period until virus elimination" and "cumulative virus production" are graphically depicted.

### <u>Supplementary Tables</u>

#### Table S1. List of compounds in the screening

| | | | | | |
|---|---|---|---|---|---|
| Abiraterone | Chlorprothixene | Favipiravir | Levodopa | Olopatadine | Rizatriptan |
| Acadesine | Cimetidine | Felodipine | Levofloxacin | Omega-3-Acid ethyl esters | Rosiglitazone |
| Acarbose | Clemastine fumarate | Fenbendazole | Levonorgestrel | Orlistat | Rutin |
| Acetohexamide | Clindamycin | Fenofibrate | Linagliptin | Oseltamivir | S-(+)-Rolipram |
| Acetylcysteine | clofibrate | Fidaxomicin | Lincomycin | Oxiconazole | Salbutamol sulfate |
| Acipimox | Clonidine | Fleroxacin | Lomustine | Oxybutynin | Simvastatin |
| Acitretin | Clotrimazole | Fluconazole | Loperamide | Oxymetazoline | Sodium butyrate |
| Acyclovir | Clozapine | Flucytosine | Lopinavir | Oxytetracycline | Sodium orthovanadate |
| Adapalene | Cortisone | Fluocinonide | Loteprednol | Ozagrel | Sorafenib |
| Adefovir Dipivoxil | Crystal violet | Flurbiprofen | Luliconazole | Paliperidone | Sotalol |
| Adenosine | Cytarabine | Fluvastatin | Manidipine | Pancuronium | Spectinomycin |
| Albendazole | Daclatasvir | Fudosteine | Maraviroc | Pazopanib HCl | Sulconazole |
| Alendronate | Daidzein | Furosemide | Masitinib | Pemafibrate | Sulfadiazine |
| Allopurinol | Dapoxetine | Gabapentin | Mesalamine | Pemetrexed | Sulfameter |
| Alogliptin | Dasatinib | Gadodiamide | Mesna | Phenylbutazone | Sulfamethoxazole |
| Amantadine | Deflazacort | Gallamine triethiodide | Mestranol | Pioglitazone | Sulfanilamide |
| Amenamevir | Delapril | Ganciclovir | Metformin | Piroxicam | Sulfisoxazole |
| Amfebutamone | Desonide | Gefitinib | Methenolone | Pitavastatin | Sulindac |
| Amiloride | Dexamethasone | Gemfibrozil | Methimazole | Potassium iodide | Tamoxifen |
| Aminocaproic acid | Dextran sulfate | Glazoprevir | Methocarbamol | Pramipexole | Taurine |
| 5-aminolevulinic acid | Dextrose | Glecaprevir | Methoxsalen | Pravastatin | Telbivudine |
| Aminophylline | Diclofenac | Gliclazide | Methylprednisolone | Praziquantel | Telmisartan |
| Amlodipine besylate | Didanosine | Glimepiride | Metolazone | Prednisolone | Teneligliptin |
| Amorolfine | Dienogest | Glipizide | Micafungin | Prilocaine | Teniposide |
| Artemether | Diethylcarbamazine | Glyburide | Miconazole | Primidone | Terbinafine |
| Atazanavir | Difluprednate | glyclopyramide | Mifepristone | Progesterone | Terguride |
| Atovaquone | Diphenhydramine | Guaifenesin | Milrinone | Pyrazinamide | Testosterone |
| Atropine | Dipyridamole | Haloperidol | Mitiglinide | Pyridostigmine | Thioguanine |
| Baloxavir marboxil | Disulfiram | Hydrochlorothiazide | Mitoxantrone | Pyrimethamine | Tofogliflozin |
| Benazepril | Divalproex sodium | Hydrocortisone | Monobenzone | Quercetin | Toremifene Citrate |
| Benserazide | Dolutegravir | Hydroxyprogesterone | Moroxydine | Quetiapine fumarate | Torsemide |
| Betamethasone | Domperidone | Hydroxyurea | Mycophenolic | Quinine | Tranilast |
| Betapar | Donepezil | Ibuprofen | Naftopidil | Racecadotril | Trelagliptin |
| Bethanechol chloride | Drospirenone | Imatinib | Nateglinide | Raltegravir | Tretinoin |
| Bezafibrate | Elbasvir | Indapamide | Nefiracetam | Ramelteon | Triamcinolone |
| Bifonazole | Elvitegravir | Indomethacin | Nelfinavir | Ramipril | Trifluridine |
| Bortezomib | Empagliflozin | Ipragliflozin | Neostigmine | Ranitidine | Trilostane |
| Bupivacaine | Emtricitabine | Ipratropium bromide | Nevirapine | Ranolazine | Tropisetron |
| Busulfan | Enalapril | Isoconazole | Niacin | Rapamycin | Ursodiol |
| Canagliflozin | Entecavir | Isoniazid | Nicomol | Relugolix | Valganciclovir |
| Captopril | Eplerenone | Isoquercetin | Nicorandil | Repaglinide | Valsartan |
| Carbamazepine | Erlotinib | Ketoprofen | Nicotinamide | Reserpine | Vandetanib |
| Carbidopa | Erythromycin | Ketorolac | Nilotinib | retinyl acetate | Vardenafil |
| Cefdinir | Estradiol | L-Glutamine | Nimodipine | Ribavirin | Vicriviroc Malate |
| Cefditoren pivoxil | Estriol | Lamivudine | Nisoldipine | Rifabutin | Vidarabine |
| Cepharanthine | Estrone | Laninamivir | Nitazoxanide | Rifampicin | Vildagliptin |
| Chenodeoxycholic acid | Ethinyl estradiol | Lanoconazole | Nitrofurazone | Rifapentine | Vitamin b12 |
| Chloramphenicol | Ethionamide | Lapatinib Ditosylate | Nizatidine | Rifaximin | Voglibose |
| Chlorothiazide | Ethyl Icosapentate | Lenalidomide | Novobiocin | Riluzole | Vorinostat |
| Chloroxine | Ezetimibe | Leuprorelin | Nystatin | Risperidone | Zalcitabine |
| Chlorpromazine | Famciclovir | Levamisole | Octreotide | Ritonavir | Zolmitriptan |

Hit compounds shown in red

**Table S2. Estimated characteristic parameters of the tested antiviral drugs**

| Drug (unit) | Class | $IC_{50}$ | $m$ |
|---|---|---|---|
| Lopinavir (μM) | RI | 1.734 | 2.992 |
| Nelfinavir (μM) | RI | 1.317 | 4.043 |
| Favipiravir (μM) | RI | $4.057 \times 10^{161}$ | $5.610 \times 10^{-3}$ |
| Chloroquine (μM) | EI | 1.313 | 1.984 |
| Cepharantine (μM) | EI | 0.991 | 3.174 |

RI, replication inhibitor; EI, entry inhibitor

$IC_{50}$, 50% inhibitory concentration

m, slope of the dose-response curve (i.e., Hill coefficient)

**Table S3. Summary of pharmacokinetic parameters of anti-SARS-CoV-2 drugs**

| Parameter name | Symbol | Unit | Nelfinavir | Cepharanthine | |
|---|---|---|---|---|---|
| | | | | i.v. | p.o. |
| Single-compartment model | | | | | |
| Maximum concentration | $C_{max}$ | μM | 2.88 | 0.278 | $5.70 \times 10^{-3}$ |
| Degradation rate | $k$ | $day^{-1}$ | 4.89 | 0.268 | 2.45 |
| Dosing schedule | | | | | |
| Initiation of treatment | $t^*$ | day | | 0.500 | |
| Dosing interval | $\tau$ | day | 0.333 | 1.00 | 1.00 |

Nelfinavir: 500 mg, TID, orally

Cepharanthine: 100mg, intravenous injection (i.v.) or 120 mg, oral administration (p.o.)

**Table S4. Estimated population parameters and initial values for SARS-CoV-2 infection**

| Parameter name | Symbol | Unit | Value |
|---|---|---|---|
| Maximum rate constant for viral replication | $\gamma$ | day$^{-1}$ | $3.16$ |
| Rate constant for virus infection | $\beta$ | (copies/ml)$^{-1}$ day$^{-1}$ | $9.77 \times 10^{-6}$ |
| Death rate of infected cells | $\delta$ | day$^{-1}$ | $0.165$ |
| Initial viral load | $V(0)$ | copies/ml | $5.64 \times 10^{3}$ |
